# ANABOLIC FAILURE, NOT CATABOLIC ACTIVATION, UNDERLIES GLYCOGEN DEPLETION IN CERVICAL CARCINOGENESIS: A MULTI-COHORT TRANSCRIPTOMIC ANALYSIS

**DOI:** 10.64898/2026.11.02.763436

**Authors:** Oyedele O. Ajayi, Efosa B Odigie, Samson O. Adesite, Theophilus O. Erameh, Anietie M. Archibong, Uche B. Eziagu

## Abstract

**Background:** Cervical epithelium loses its glycogen as disease grade rises, an observation old enough to underpin iodine-based visual screening, but which arm of glycogen metabolism accounts for the loss has not been established. The expectation from cancer metabolism is catabolic: that stored glycogen is mobilised and glycolysis activated.

**Methods:** We analyzed 238 specimens from four publicly available cohorts across three sample types: 128 laser-capture-microdissected epithelial samples, 35 bulk-tissue samples, and 75 exfoliated cytology samples in two sets (44 and 31; GSE27678). We prespecified a 33-gene panel comprising four modules: glycogen synthesis, glycogen degradation, glycolysis, and squamous differentiation. For each specimen, we calculated module scores using single-sample gene set enrichment. Differential expression was assessed with limma, pathway enrichment with fgsea, and classification performance with bootstrap confidence intervals for ROC curves. We documented all analytical decisions and their rationale.

**Results:** Synthesis and differentiation fell with grade in all four cohorts, while degradation and glycolysis stayed at their normal-tissue baseline. All seven differentiation genes were reduced in carcinoma and all seven occupied the top 5.1% of the ranked transcriptome, with CRNN the most down-regulated of 21,755 genes. Synthesis failed selectively, through GYS2 and PPP1R3C only, with the ubiquitous synthase GYS1 unchanged. The glycolytic module separated from normal epithelium at no grade (p 0.45 to 0.62), classified CIN3 or worse at an area under the curve of 0.569 with an interval spanning chance, and showed no enrichment in carcinoma. Synthesis separated from normal first, at CIN2, and differentiation at CIN3. The composite index tracked grade in the discovery cohort (Spearman rho 0.652, *p* 7.9×10^-17^), but offered no discriminatory advantage over its differentiation component alone (0.832 against 0.832).

**Conclusions:** Glycogen depletion in cervical carcinogenesis reflects failure of a tissue-restricted synthetic axis, GYS2 and PPP1R3C, within a collapsing squamous differentiation programme, not catabolic activation. Measuring all four candidate mechanisms on one scale in the same specimen is what allows the catabolic explanation to be excluded rather than merely left untested. The pattern holds across two array chemistries and three specimen preparations, including the exfoliated cytology that screening already collects, and the synthesis and differentiation modules together are the molecular counterpart of what iodine staining reports. These are transcript-level findings and define no clinical threshold.

## Introduction

Cervical cancer remains among the most common cancers in women worldwide and is a leading cause of cancer-related mortality in low- and middle-income countries. Its burden is disproportionately concentrated in these settings, where limited infrastructure often restricts access to routine cytology- or HPV-based screening[1– 5]. In these settings, visual inspection methods carry the front-line screening load, recommended by the World Health Organization and deployed in national programmes across sub-Saharan Africa, South Asia and Latin America [6–9]. Both rest on the same underlying tissue change. Visual inspection with Lugol’s iodine (VILI) reads it directly: iodine binds the glycogen of mature squamous epithelium and stains it mahogany brown, while glycogen-depleted lesions fail to take up stain. Visual inspection with acetic acid (VIA) reads an epithelium whose glycogen has already been lost, the acetowhitening response itself arising from reversible coagulation of nuclear and cytoskeletal protein against a raised nuclear-to-cytoplasmic ratio [10–12]. Both are read by the naked eye, require no laboratory infrastructure, and underpin the screen-and-treat approach credited with measurable reductions in cervical cancer incidence in resource-constrained populations . Glycogen depletion is therefore the tissue state on which this entire class of screening depends, yet it has not been resolved at the level of gene-expression mechanism.

That depletion has been demonstrable histologically since Schiller introduced iodine staining in 1933 and is routinely confirmed by periodic acid-Schiff staining[13–16]. Normal squamous cervical epithelium accumulates glycogen in its intermediate and superficial layers as a property of maturation, and dysplastic epithelium loses most of it[17,18]. What remains unclear is the molecular driver. Glycogen loss could in principle arise from upregulated catabolism to fuel proliferation[19,20], from downregulated synthesis machinery, from a Warburg-style shift that diverts glucose away from storage[21–23], or from collapse of the squamous differentiation programme that distinguishes glycogen-rich upper-layer cells from undifferentiated basal cells [24–27]. These four mechanisms make opposite predictions about which genes move and in which direction, and they are not distinguishable by histochemistry, which reports the presence of glycogen but not the reason for its absence. No published transcriptomic study has quantified all four arms in parallel across a multi-cohort design with controlled platform and specimen variation.

Resolving this mechanistic question matters for three reasons. Therapeutically, the dominant published framing has positioned cervical metabolic reprogramming as Warburg-driven [28,29], an interpretation that has shaped investment in glycolytic-axis drug development and biomarker discovery [23,28–31]. If the primary mechanism is anabolic failure layered onto differentiation loss rather than catabolic acceleration, the therapeutic logic shifts away from glycolytic inhibitors and toward agents that engage the squamous differentiation program [32–35] or restore protein phosphatase 1 targeting of glycogen synthase [20,36–39] . Diagnostically, whether the signature is detectable in exfoliated cytology, the specimen type visual screening already collects, and whether it behaves consistently across array chemistries, are prerequisites for any molecular adjunct and remain unestablished. For population health, the biology underlying visual screening deserves direct molecular characterisation; a class of methods that informs clinical decisions for tens of millions of women annually should rest on more than histological precedent [26,40,41] . A multi-cohort, multi-platform, multi-specimen test of the underlying transcriptomic biology is therefore needed.

We tested whether a curated gene-expression signature covering the four mechanistic arms of cervical glycogen biology, namely synthesis, degradation, Warburg-axis glycolysis and squamous differentiation, tracks disease grade and behaves consistently across array platform and specimen preparation [20,26,42–44]. Three questions guide this study. First, does the signature change progressively from normal epithelium through CIN1, CIN2 and CIN3 to invasive cancer, and at what grade does each arm first differ from normal? Second, does a composite index combining all four arms discriminate disease better than the individual arms, or is the signal driven by one arm? Third, are findings from laser-microdissected epithelium consistent in bulk tissue and exfoliated cytology, the specimen type a deployable test would use?

Three features strengthen this test. First, all four arms are measured and reported in parallel, so an unchanged arm remains an informative result. That matters most where the prevailing framing predicts movement, which is the catabolic side. Second, platform and specimen preparation vary independently: the same Affymetrix array covers both microdissected epithelium and exfoliated cytology, while a single cytology series spans two array chemistries. This design helps separate platform effects from specimen effects. Third, we document analytical decisions and their rationales, including those made after data inspection, and report outlier sensitivity, leave-one-gene-out and module dropout analyses, together with instances where the composite index does not outperform its components. Negative results are meaningful only when readers can assess how they were obtained.

## Methods

### 2.1 Cohorts and specimens

Four cohorts were retrieved from the Gene Expression Omnibus with GEOquery. GSE63514 (GPL570, Affymetrix HG-U133 Plus 2.0) provided 128 laser-capture-microdissected cervical epithelial specimens graded normal, CIN1, CIN2, CIN3 and invasive carcinoma. GSE138080 (GPL4133, Agilent-014850 Whole Human Genome 4×44K, single channel) provided 35 bulk tissue specimens graded normal, high-grade cervical intraepithelial neoplasia (HGCIN, equivalent to CIN2/3) and squamous cell carcinoma, following the histological labels used by the series itself; five specimens carry grade 2-3 neoplasia in their characteristics field rather than HGCIN in the title and were assigned on that basis. GSE27678 contains two sub-platforms, which were separated by querying the platform annotation of each returned expression set rather than by list position: GPL571 (HG-U133A 2.0) provided 44 exfoliated cytology specimens graded normal, LSIL and HSIL, and GPL570 provided 33 exfoliated cytology specimens. Two of the GPL570 specimens are immortalised cell lines, MS751 (GSM685543) and CaSki (GSM685549); both were excluded, since a cell line has no stratified epithelium and therefore cannot carry the differentiation gradient the panel measures, leaving 31 specimens. These three preparations, microdissected epithelium, bulk tissue and exfoliated cytology, differ in cell-type composition and are reported separately throughout.

One GPL570 specimen, GSM685524, carries conflicting deposited metadata: its tissue and type fields record normal uterine cervix, while its source field records squamous cell carcinoma. It was retained as normal on the agreement of two of three fields, and because retention is the conservative choice for the reported estimate. A post hoc analysis excluding it is reported separately and is not combined with any pre-specified analysis.

### 2.2 Processing and quality control

Expression matrices were checked for scale before any analysis. GSE138080 was deposited on a linear scale (median 110, maximum 650,658) and was log2 transformed. Per-array median intensities after transformation spanned 0.19 log2 units, so no between-array normalisation was applied; the alternative quantile-normalised matrix was retained and gave the same conclusions. The remaining three matrices were already log2 scaled.

Probes were collapsed to gene symbols by MaxMean selection, retaining for each gene the probe with the highest mean expression across samples. This replaces averaging across replicate probes. Probe failure from poor hybridisation kinetics, non-functional splice targets or 3-prime bias is common, whereas cross-hybridisation producing a spuriously high signal is rare, so the highest-mean probe is the better single estimate of the dominant transcript; averaging instead draws a gene toward background wherever its probes differ widely in intensity. The cost of this choice is the loss of variance reduction from averaging genuine replicate probes, which is accepted in exchange for removing a systematic bias. Results under both rules are reported in **Supplementary Table S11b**, and the probe selected for each panel gene, with the number of probes available and the spread of their mean expression, in **Supplementary Table S11a**.

Quality control comprised inter-sample expression boxplots, density curves, a sample-to-sample correlation matrix, and principal component analysis on the 2,000 most variable genes of the collapsed matrix. A second principal component analysis on all genes was computed as a concordance check and is reported in the run log rather than as a figure. Two outlier criteria were pre-specified: distance from the group centroid in the first two principal components, z-scored within group, exceeding 3; and mean within-group correlation, z-scored within group, below -3. No specimen met the principal component criterion, the largest centroid z being 2.30. Four specimens met the correlation criterion. The threshold was not adjusted after inspection

### 2.3 Gene panel

Thirty-three genes were assigned in advance to four modules, in the fixed order Synthesis, Degradation, Warburg, Differentiation, used throughout the figures and tables. Synthesis comprises GYS1, GYS2, GBE1, UGP2, PGM1, PPP1R3A, PPP1R3B, PPP1R3C and PPP1R3D; Degradation comprises PYGB, PYGL, PYGM, AGL, PHKA1, PHKA2, PHKB, PHKG1 and PHKG2; Warburg comprises HK2, PFKL, PKM, LDHA, ENO1, SLC2A1, SLC2A3 and G6PD; Differentiation comprises KRT4, KRT13, IVL, SPRR1A, TGM3, CRNN and ESR1. Genes were selected for a documented role in cervical glycogen biology or the wider squamous metabolic literature, detectable expression in keratinised epithelium, and presence on at least three of the four arrays.

Cohort-specific gene sets were formed by intersecting the locked panel with available symbols. For GSE138080, probes carrying an Entrez identifier but no gene symbol were mapped through org.Hs.eg.db in an attempt to recover missing panel members; none of the five absent genes was recovered, and this negative result is reported. The panel was not modified after inspection of the data, including after two members were found to be expressed at background in the discovery cohort.

### 2.4 Differential expression

Differential expression used limma with a cell-means design, model.matrix with no intercept on the grade factor, and seven contrasts: CIN1, CIN2, CIN3 and carcinoma each against normal, and CIN2 against CIN1, CIN3 against CIN2 and carcinoma against CIN3. Fits proceeded through lmFit, contrasts.fit and eBayes, with topTable extracted for all genes without sorting and Benjamini-Hochberg adjustment applied within contrast. Significance was defined as adjusted p below 0.05; where a fold-change filter is also applied it is stated explicitly as exceeding one log2 unit.

The analysis was repeated on three specimen sets, which are reported separately and never pooled: the full cohort; the cohort excluding the four specimens meeting the pre-specified outlier criteria; and the cohort excluding the five specimens named in an earlier version of this work, which met neither stated criterion and are reported only for continuity. The full transcriptome-wide results for the pre-specified sensitivity set are given in **Supplementary Table S3** and for the continuity set in **Supplementary Table S3b**.

### 2.5 Single-sample enrichment and the composite index

Module scores were computed per sample with GSVA, using ssgseaParam on the collapsed gene-level matrix with the cohort-specific gene sets and default normalisation. Two forms of a composite index were defined in advance of analysis:

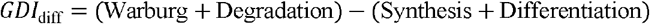

and the corresponding ratio of the same two sums. The difference form was pre-specified as primary. The ratio form is undefined where its denominator approaches zero and changes sign where the denominator becomes negative, which occurs in three discovery specimens; it is reported for completeness and carries no conclusion.

ssGSEA normalises against the full sample set supplied to it, so module scores and both index forms are internal to a cohort. Absolute values are not comparable between cohorts and no transferable threshold is implied. Every subset analysis recomputed enrichment from the subset expression matrix rather than taking a subset of previously computed scores.

### 2.6 Trend and discrimination

Association with ordinal grade was tested by Spearman correlation with the asymptotic approximation and by Kruskal-Wallis, in all four cohorts and for both index forms; the H statistic and its degrees of freedom are reported alongside p. GSE27678 GPL570 contains two groups, so its Spearman coefficient is a two-group comparison and is described as such rather than as a trend.

Discrimination used pROC with thresholds fixed in advance: CIN2 or worse, CIN3 or worse and carcinoma in the discovery cohort; HGCIN or worse and SCC in GSE138080; LSIL or worse and HSIL in GPL571; and carcinoma in GPL570. Confidence intervals came from 2,000 stratified bootstrap resamples. For each estimate the number of positive and negative specimens and the number of discordant pairs are reported, because several estimates rest on small groups.

The discovery cohort is the primary analysis. GSE138080 and GPL571 are replication cohorts in which the trend statistic, which uses every specimen, is the primary measure and each area under the curve is reported with its interval and both group sizes. GPL570 carries three normal specimens and is reported as a binary concordance check rather than as a validation.

### 2.7 Robustness

Three analyses tested dependence on individual specimens and genes. Outlier sensitivity recomputed enrichment, the index and all three discovery estimates after removing each exclusion set in turn, comparing the shift in each estimate against the half-width of its bootstrap interval (**Supplementary Table S9f)**. Leave-one-gene-out removed each of the 33 genes from its module, recomputed enrichment and the index from the reduced set, and re-estimated the CIN3-or-worse area under the curve; any gene whose removal would empty a module would have been skipped, and none did. Module dropout tested each module alone as a classifier, inverting Synthesis and Differentiation so that low expression yields a high disease score, and compared each against the composite. Reduced index forms were compared on the same thresholds to test whether each module earns its place.

### 2.8 Pathway enrichment

Genes were ranked by the signed significance of the carcinoma versus normal contrast, as the sign of the log2 fold change multiplied by the negative log10 of the nominal p value with a floor added to avoid infinities, using the nominal rather than the adjusted p. Non-finite and exactly zero values were dropped and the list sorted in decreasing order. The four modules were tested against this list with fgsea, minimum set size 5, maximum 50, and 10,000 simple permutations.

The leading edge returned for a module contains only members moving in the direction of its enrichment score, so it is not a list of the module’s responsive genes. Significant members are therefore also tabulated by direction, and the rank position of every panel gene in the full list is reported.

### 2.9 Software and reproducibility

Analyses ran under R 4.5.3 with Bioconductor 3.22 on Windows, using GEOquery 2.78.0, limma 3.66.0, GSVA 2.4.9, fgsea 1.36.2, pROC 1.19.0.1, together with Biobase, AnnotationDbi, org.Hs.eg.db, pheatmap, ggplot2, patchwork and dplyr. All figures are ggplot2 output from the deposited scripts. Analysis code was developed in R with assistance from an AI assistant (Claude, Anthropic), used interactively for script drafting, debugging and documentation. All analytical decisions, parameter choices and interpretations were made by the authors. Every reported value is reproduced by the deposited code, and a verification script checks thirty reported quantities against the manuscript text. A decision record documenting each analytical choice, including corrections made during review, is provided as **Supplementary Table S12** and in the repository.

## Result

Glycogen depletion across cervical disease grade is driven by loss of glycogen synthesis and squamous differentiation, with no detectable contribution from catabolic or glycolytic programmes.

### 3.1 Study populations and panel coverage

Two hundred and forty specimens were available across the four cohorts. Two were excluded, both immortalised cell lines in the GSE27678 GPL570 set, leaving 238 for analysis. The analysed specimens comprise 49 normal, 25 low-grade lesions, 22 CIN2, 76 high-grade lesions and 66 invasive carcinomas, distributed across three specimen preparations: 128 laser-capture-microdissected epithelia, 35 bulk tissues and 75 exfoliated cytology specimens. Cohort composition, platform and preparation are given in **Table 1**, and quality-control diagnostics for each cohort in **Figure S1**.

**Table 1.** Cohorts, platforms, specimen preparations and composition. Panel coverage gives the number of the 33 panel genes present on each platform. Five genes are absent from the GPL4133 design: GYS2, PPP1R3A, PHKG1, SLC2A3 and TGM3. PPP1R3B is absent from GPL571. The GSE27678 GPL570 set is given after exclusion of two immortalised cell lines, MS751 and CaSki.

| **Role** | **Accession** | **Platform** | **Preparation** | **Composition** | **n** | **Genes** | **Panel coverage** |
|---|---|---|---|---|---|---|---|
| Discovery | GSE63514 | Affymetrix HG-U133 Plus 2.0 (GPL570) | Laser-capture-microdissected epithelium | Normal 24, CIN1 14, CIN2 22, CIN3 40, Cancer 28 | 128 | 21755 | 33 of 33 |
| Replication 1 | GSE138080 | Agilent-014850 Whole Human Genome 4x44K (GPL4133) | Bulk tissue | Normal 10, HGCIN 15, SCC 10 | 35 | 13609 | 28 of 33 |
| Replication 2 | GSE27678 | Affymetrix HG-U133A 2.0 (GPL571) | Exfoliated cytology | Normal 12, LSIL 11, HSIL 21 | 44 | 13101 | 32 of 33 |
| Concordance check | GSE27678 | Affymetrix HG-U133 Plus 2.0 (GPL570) | Exfoliated cytology | Normal 3, Cancer 28 | 31 | 21755 | 33 of 33 |

Panel coverage was 33 of 33 in the two GPL570 cohorts, 32 of 33 in GPL571, which lacks PPP1R3B, and 28 of 33 in GSE138080 (Table 1). The five genes absent there, GYS2, PPP1R3A, PHKG1, SLC2A3 and TGM3, are not represented in the GPL4133 annotation, and mapping unannotated probes through Entrez identifiers recovered none of them. Three panel genes are not detected above array background in normal cervical epithelium: PYGM (mean log2 1.90), PHKG1 (1.67) and PPP1R3A (3.02). All three are the skeletal-muscle isoforms of their enzymes, while the liver and brain isoforms of the same enzymes are well expressed, so their absence is the expected behaviour of squamous epithelium. All three were retained, and their contribution is quantified in Section 3.7.

### 3.2 Differential expression across disease grade

Transcriptome-wide differential expression across 21,755 genes separated the grades sharply but unevenly (**Figure S2, Table S2**). Against normal epithelium, CIN1 yielded 1 significant gene at FDR below 0.05, CIN2 yielded 311, CIN3 yielded 5,423 and carcinoma yielded 8,555. Between consecutive grades, CIN2 versus CIN1 yielded no significant genes at any threshold, with a minimum FDR of 0.995, while CIN3 versus CIN2 yielded 1,048 and carcinoma versus CIN3 yielded 1,808.

Among the 33 panel genes in the carcinoma versus normal contrast, 13 reached FDR below 0.05 and 11 of these also exceeded one log2 unit of change (**Table S5**). Every significant Differentiation gene fell, and all seven members of that module were significant: CRNN at -9.509 (FDR 3.2e-12), KRT4 at -5.460 (1.5e-06), KRT13 at -4.670 (8.5e-06), IVL at -4.244 (9.5e-08), ESR1 at -4.121 (2.3e-07), TGM3 at -3.962 (4.5e-05) and SPRR1A at -3.533 (1.6e-04). CRNN occupies the final position in the ranked transcriptome, making cornulin the most strongly down-regulated gene of the 21,755 tested **(Figure 1A)**. Expression of all 33 panel genes across the five grades is shown in **Figure 1B**, and per-gene values with platform availability in **Table 2**.

**Table 2.** The 33-gene panel by module, with platform availability and behaviour in the discovery cohort. The 33-gene panel and its behaviour in the discovery cohort. Genes appear in the locked module order. Log2 fold change, false discovery rate and rank are for the carcinoma versus normal contrast across 21,755 genes, where rank 1 is the most up-regulated. Mean log2 expression is given for the normal group, because detectability is a property of the tissue before disease rather than after it. The Notes column flags the five genes absent from the Agilent GPL4133 array and the three expressed at array background in normal epithelium. Per-cohort availability and mean expression across all four cohorts are given in Supplementary Table S1.

| Module | Gene | log2FC | FDR | Rank | Mean log2 expression, normal | Notes |
|---|---|---|---|---|---|---|
| Synthesis | GYS1 | -0.447 | 0.297 | 14127 | 7.18 |  |
| Synthesis | GYS2 | -3.742 | 2.20E-06 | 21163 | 6.09 | absent from GPL4133 |
| Synthesis | GBE1 | 0.201 | 0.589 | 8360 | 10.84 |  |
| Synthesis | UGP2 | 0.073 | 0.84 | 9500 | 12.2 |  |
| Synthesis | PGM1 | -0.296 | 0.417 | 13279 | 10.54 |  |
| Synthesis | PPP1R3A | -0.107 | 0.201 | 15047 | 3.02 | absent from GPL4133; below array background |
| Synthesis | PPP1R3B | -0.363 | 0.556 | 12425 | 8.96 | absent from GPL571 |
| Synthesis | PPP1R3C | -5.124 | 4.12E-06 | 21109 | 11.89 |  |
| Synthesis | PPP1R3D | 0.211 | 0.445 | 7634 | 7.83 |  |
| Degradation | PYGB | -0.732 | 0.004 | 19840 | 7.81 |  |
| Degradation | PYGL | -0.337 | 0.401 | 13368 | 11.09 |  |
| Degradation | PYGM | -0.011 | 0.407 | 13340 | 1.9 | below array background |
| Degradation | AGL | 0.949 | 0.009 | 2944 | 11.56 |  |
| Degradation | PHKA1 | 0.786 | 0.108 | 5166 | 6.08 |  |
| Degradation | PHKA2 | 0.396 | 0.167 | 5737 | 7.95 |  |
| Degradation | PHKB | -0.247 | 0.426 | 13204 | 10.37 |  |
| Degradation | PHKG1 | -0.046 | 0.567 | 12357 | 1.67 | absent from GPL4133; below array background |
| Degradation | PHKG2 | 1.265 | 0.004 | 2411 | 5.69 |  |
| Warburg | HK2 | 0.44 | 0.233 | 6273 | 11.94 |  |
| Warburg | PFKL | -0.208 | 0.52 | 12620 | 9.42 |  |
| Warburg | PKM | -0.405 | 0.488 | 12790 | 11.14 |  |
| Warburg | LDHA | 0.015 | 0.94 | 9953 | 14.72 |  |
| Warburg | ENO1 | 0.322 | 0.622 | 8501 | 13.11 |  |
| Warburg | SLC2A1 | 0.248 | 0.472 | 7758 | 10.09 |  |
| Warburg | SLC2A3 | 1.715 | 0.005 | 2553 | 6.18 | absent from GPL4133 |
| Warburg | G6PD | 0.138 | 0.794 | 9301 | 7.61 |  |
| Differentiation | KRT4 | -5.46 | 1.48E-06 | 21203 | 13.9 |  |
| Differentiation | KRT13 | -4.67 | 8.45E-06 | 21028 | 14.43 |  |
| Differentiation | IVL | -4.244 | 9.52E-08 | 21469 | 11.84 |  |
| Differentiation | SPRR1A | -3.533 | 1.61E-04 | 20640 | 13.96 |  |
| Differentiation | TGM3 | -3.962 | 4.48E-06 | 20821 | 6.65 | absent from GPL4133 |
| Differentiation | CRNN | -9.509 | 3.18E-12 | 21747 | 12.97 |  |
| Differentiation | ESR1 | -4.121 | 2.28E-07 | 21372 | 12.34 |  |

**Figure 1.**
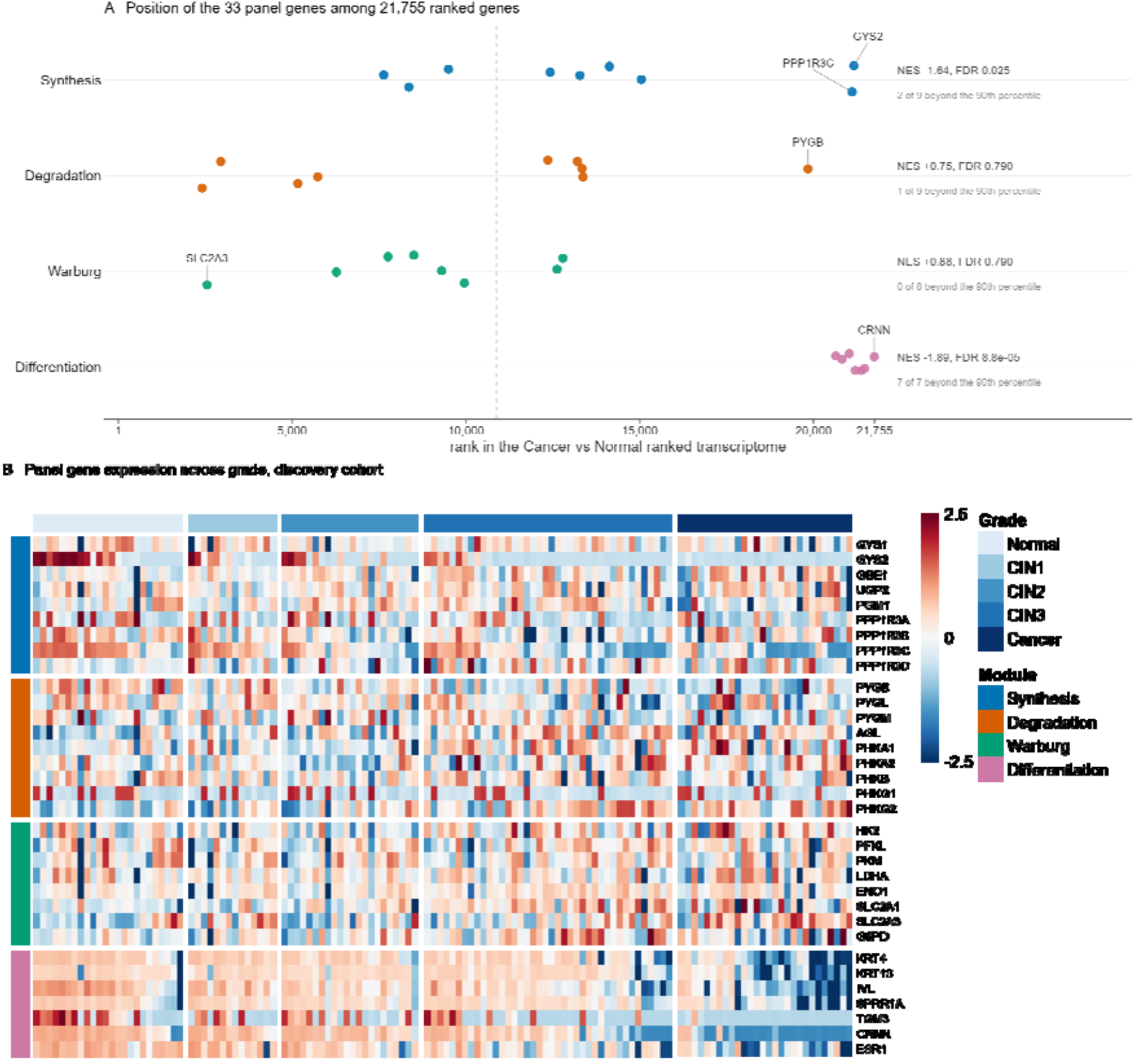
Behaviour of the 33 panel genes in the discovery cohort. (A) Position of each panel gene in the Cancer versus Normal ranked transcriptome, where rank 1 is the most up-regulated and rank 21,755 the most down tail, Synthesis contributes two extreme members and seven near the centre, and no Warburg member reaches either tail. (B) Expression of the same 33 genes across all 128 specimens, ordered by grade and, within grade, by ascending index value. Rows follow the locked module order: Synthesis 9, Degradation 9, Warburg 8, Differentiation 7. Values are z-scores computed within gene and clipped at plus or minus 2.5, the 98th percentile of absolute z.

The Synthesis module behaved differently. Only two of its nine members moved: PPP1R3C at -5.124 (4.1e-06) and GYS2 at -3.742 (2.2e-06). The ubiquitous synthase GYS1 did not change (-0.447, FDR 0.297), nor did GBE1, UGP2, PGM1 or the remaining regulatory subunits. Loss of glycogen synthetic capacity in this tissue is therefore selective for the hepatic-type synthase and the PTG glycogen-targeting subunit rather than general across the pathway.

The Degradation module moved in both directions. PHKG2 rose by 1.265 (3.6e-03) and AGL by 0.949 (9.4e-03), while PYGB fell by -0.732 (4.3e-03); the remaining six members did not change. In the Warburg module only SLC2A3 reached significance, rising by 1.715 (4.7e-03); SLC2A1, HK2, ENO1, PKM, PFKL, LDHA and G6PD did not change. Module mean log2 fold changes were -1.066 for Synthesis, +0.225 for Degradation, +0.283 for Warburg and -5.071 for Differentiation.

### 3.3 The CIN1 to CIN2 transition

No gene in the transcriptome distinguishes CIN2 from CIN1 in this cohort. The minimum FDR across all 21,755 genes is 0.995 (**Table S2**). Four panel genes move by more than 0.8 log2 units at this transition, PPP1R3C at -2.907, SLC2A3 at -1.533, KRT4 at -1.084 and PPP1R3B at -0.835, but each carries an FDR of 1.000 (**Table S5**). No claim of an early gene-level metabolic event is supportable here.

The transition is detectable at module level. Comparing each grade against normal in the discovery cohort, the Synthesis module separates first, at CIN2 (median 0.2152 against 0.2902, Mann-Whitney p = 0.024). Differentiation does not separate until CIN3 (0.4140 against 0.5171, p = 2.19e-04), having failed to reach significance at CIN2 (p = 0.088). The composite index follows Differentiation rather than Synthesis, separating from normal only at CIN3 (p = 1.19e-04). The Degradation module crosses the threshold at CIN2 (p = 0.011) and falls back below it at CIN3 and in carcinoma, an excursion rather than a trend; **Figure 2A** shows it rising and returning while Synthesis and Differentiation continue to climb. The Warburg module does not separate from normal at any grade, with p values between 0.45 and 0.62. These comparisons are unadjusted across modules and grades, so they establish ordering rather than individual significance (**Figure 2A, Table S4**).

**Figure 2.**
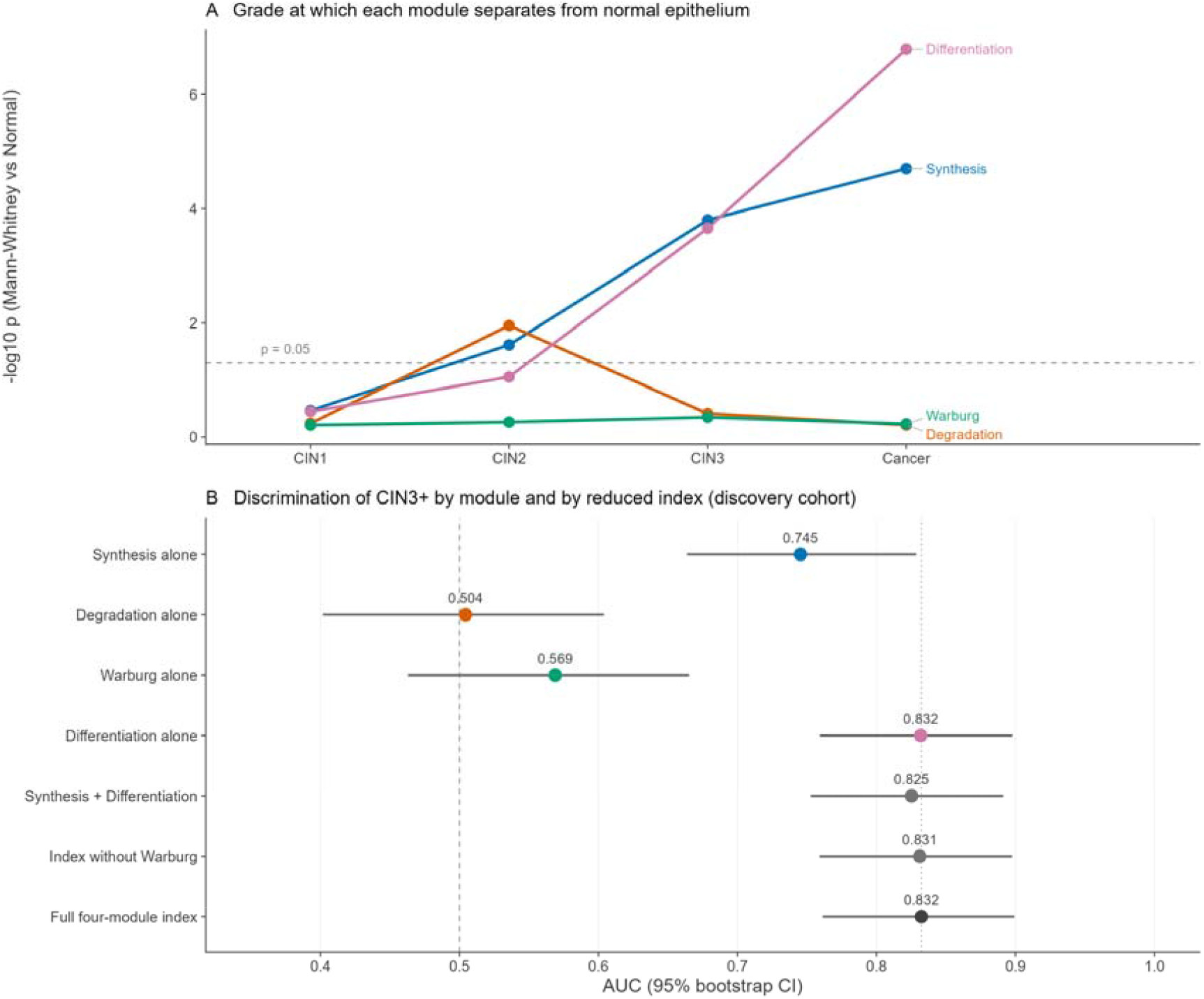
Location of the signal within the panel. (A) Grade at which each module separates from normal epithelium in the discovery cohort, plotted as the negative log of the Mann-Whitney p value against normal for each grade. The dashed line marks p = 0.05. Synthesis crosses first, at CIN2, and Differentiation at CIN3; Degradation crosses at CIN2 and returns below the line thereafter; Warburg does not cross at any grade. These comparisons are unadjusted across modules and grades and establish ordering rather than individual significance. (B) Discrimination of CIN3 or worse by each module alone and by reduced forms of the composite index, with 95% confidence intervals from 2,000 stratified bootstrap resamples. Synthesis and Differentiation are inverted so that low expression yields a high disease score. The dotted line marks the full four-module index. Differentiation alone is equivalent to the composite, and the intervals for Degradation and Warburg both include 0.5.

The ordering is the substantive finding: glycogen synthetic capacity declines before the squamous differentiation programme fails, and both precede any detectable change in catabolic or glycolytic gene expression.

### 3.4 Module trajectories across grade and cohort

Plotting each module as its change from the cohort’s own normal median shows the same pattern in all four cohorts (**Figure 3, Table S7**). Synthesis and Differentiation fall with grade; Degradation and Warburg stay close to baseline throughout, their largest excursion in any cohort being 0.076. In the discovery cohort the Synthesis median moves from 0.2902 in normal epithelium to 0.1753 in carcinoma, and Differentiation from 0.5171 to 0.1269. Over the same interval Degradation moves from 0.1754 to 0.1726 and Warburg from 0.3970 to 0.4017, changes of 0.003 and 0.005 against a Differentiation change of 0.390. In the GPL570 concordance check both catabolic modules rise rather than fall, by 0.049 and 0.060, the opposite direction to depletion, against a Differentiation fall of 0.417 in the same panel

**Figure 3.**
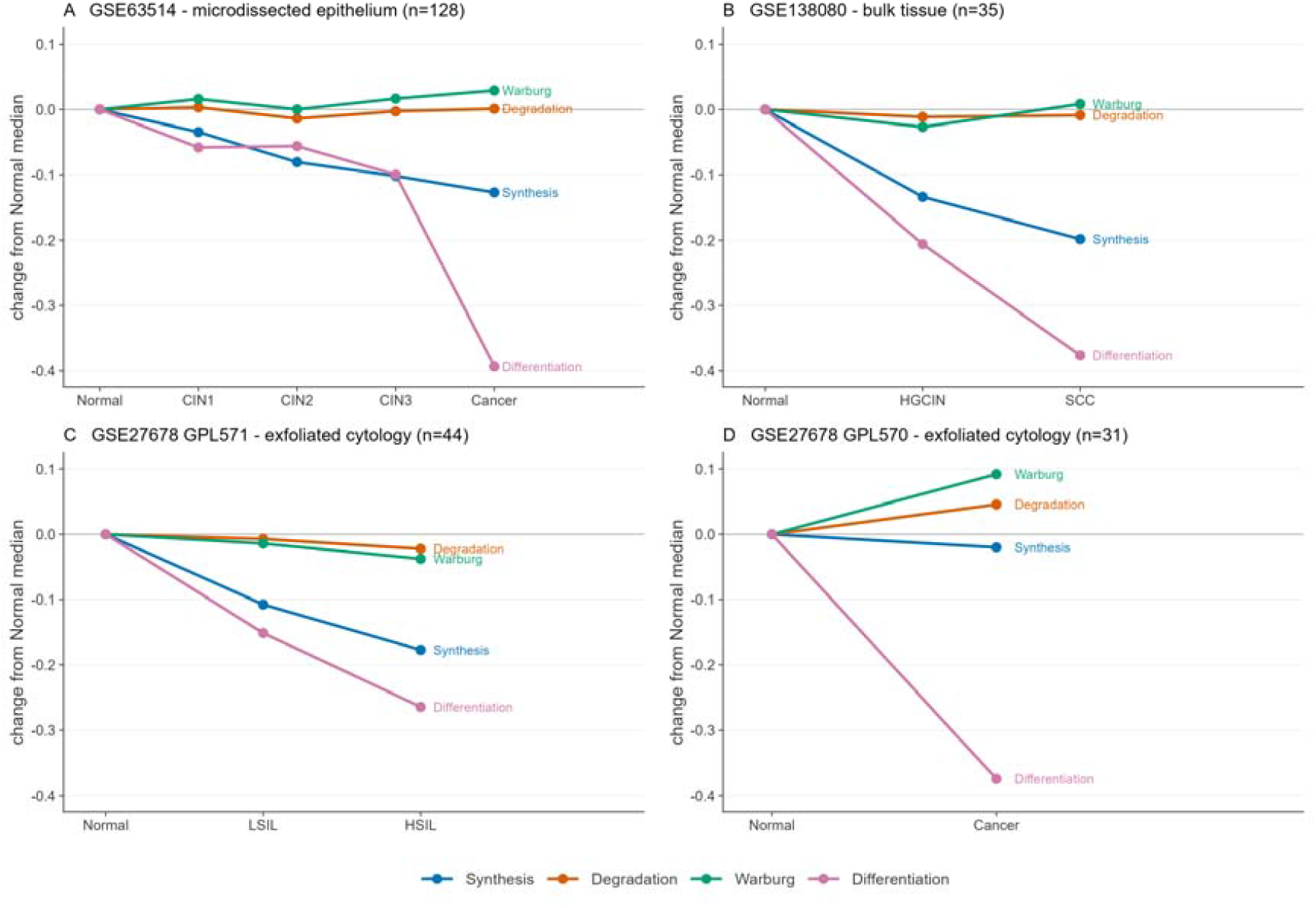
Module score trajectories across disease grade in four cohorts. Each panel plots the change in median single-sample enrichment score from that cohort’s own normal median, for each of the four modules, on a shared vertical scale. Enrichment scores are normalised within a sample set, so absolute values are internal to a cohort and not comparable between panels, whereas differences from the cohort’s own normal group are. Specimen preparation and sample size are given in each panel title. Synthesis and Differentiation fall with grade in all four cohorts, while Degradation and Warburg stay within 0.08 of the normal baseline in every cohort; in panel D, a two-group comparison against three normal specimens, both rise slightly. Colours are the Okabe-Ito colourblind-safe set, and every line is directly labelled.

The magnitude of the Differentiation decline varies by specimen preparation, falling by 0.39 in microdissected epithelium, 0.37 in bulk tissue, 0.42 in the GPL570 cytology set and 0.23 in the GPL571 cytology set. Absolute ssGSEA scores are not comparable between cohorts, because the method normalises within each sample set; only changes from a cohort’s own normal group are interpretable, and no transferable cut-point exists.

The composite index, GDI_diff, rises monotonically across the five discovery grades (-0.236, -0.202, -0.170, - 0.048, +0.245). Monotonicity is not claimed. Under the alternative probe-collapse rule the normal to CIN1 step reverses, so the ordering at that step depends on an analytical choice rather than on the data. That step is in any case not significant under either rule (Mann-Whitney p = 0.44; bootstrap 95% confidence interval on the median difference -0.074 to +0.141, with CIN1 exceeding normal in 78.2% of 2,000 resamples). The Spearman trend, reported below, is the evidence for grade-dependence.

The ratio form of the index is reported in **Table S7 and S8** for completeness but carries no conclusion. Its denominator crosses zero in three discovery samples (GSM1551331, GSM1551419, GSM1551428), producing ratios of -3.150, -3.496 and -3.005 and a cohort range from -3.496 to +7.423. Within carcinoma its standard deviation is 8.9 times that of the difference form.The instability is larger still in the smaller cohorts: in the GPL570 carcinoma group the ratio runs from -57.26 to +16.14 with a standard deviation of 11.68, against 0.247 for the difference form in the same specimens.

### 3.5 Trend and discrimination

The index tracks ordinal grade in every cohort (**Table 3**; both index forms in **Table S4**). In the discovery cohort Spearman rho is 0.652 (p = 7.9e-17) with Kruskal-Wallis H = 57.14 on 4 degrees of freedom (p = 1.2e-11). In GSE138080 rho is 0.719 (p = 1.2e-06, H = 18.19, 2 df), and in GSE27678 GPL571 rho is 0.575 (p = 4.5e-05, H = 14.29, 2 df). The difference form outperformed the ratio form on every statistic in every cohort.

**Table 3.**
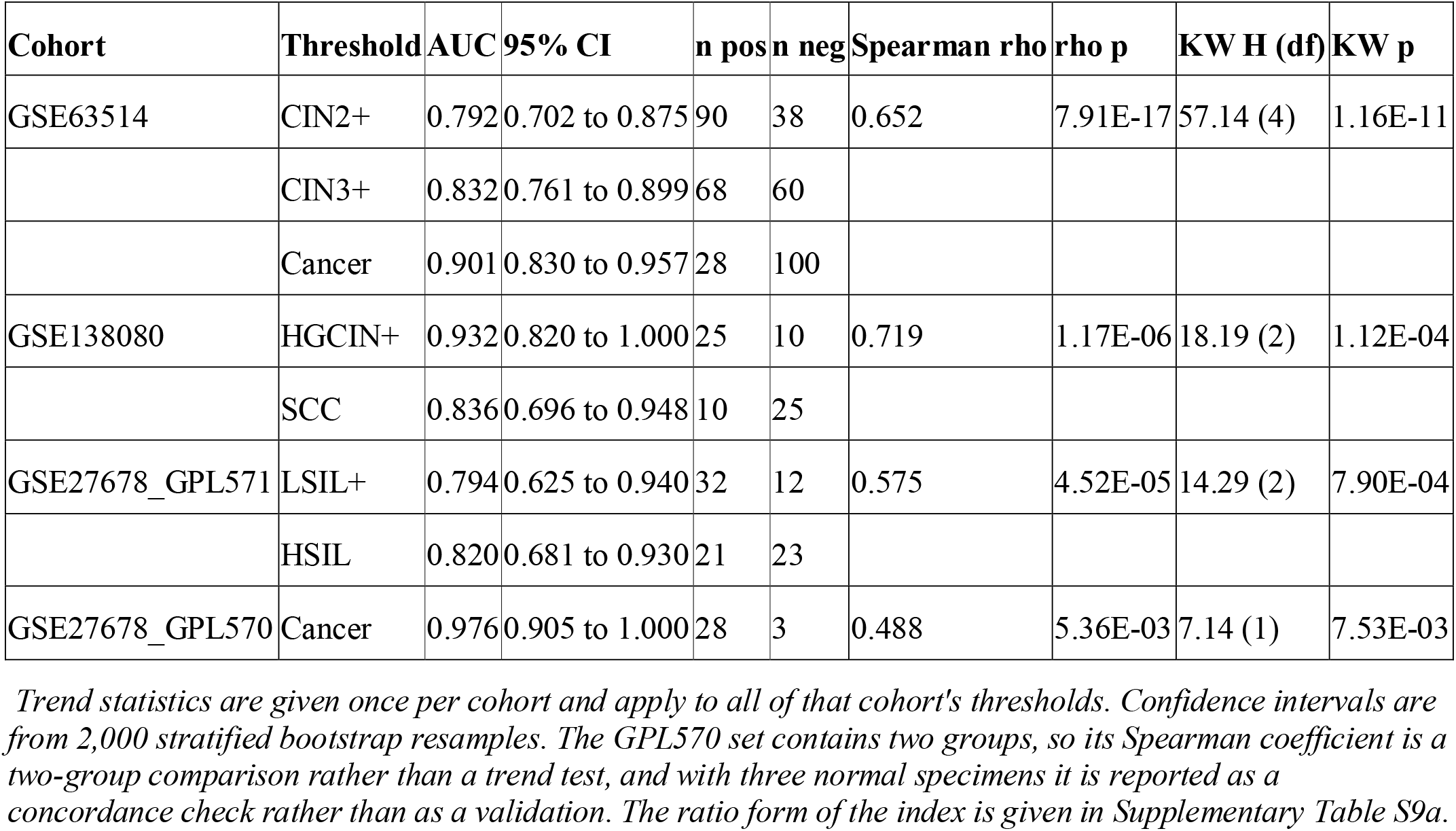
Trend statistics and discrimination for the composite index in its pre-specified difference form.

Discrimination in the discovery cohort gave areas under the curve of 0.792 (95% CI 0.702 to 0.875) for CIN2 or worse, 0.832 (0.761 to 0.899) for CIN3 or worse, and 0.901 (0.830 to 0.957) for carcinoma (**Figure 4, Table 3**). These are the only estimates in the study with at least 38 specimens on each side of the threshold.

**Figure 4.**
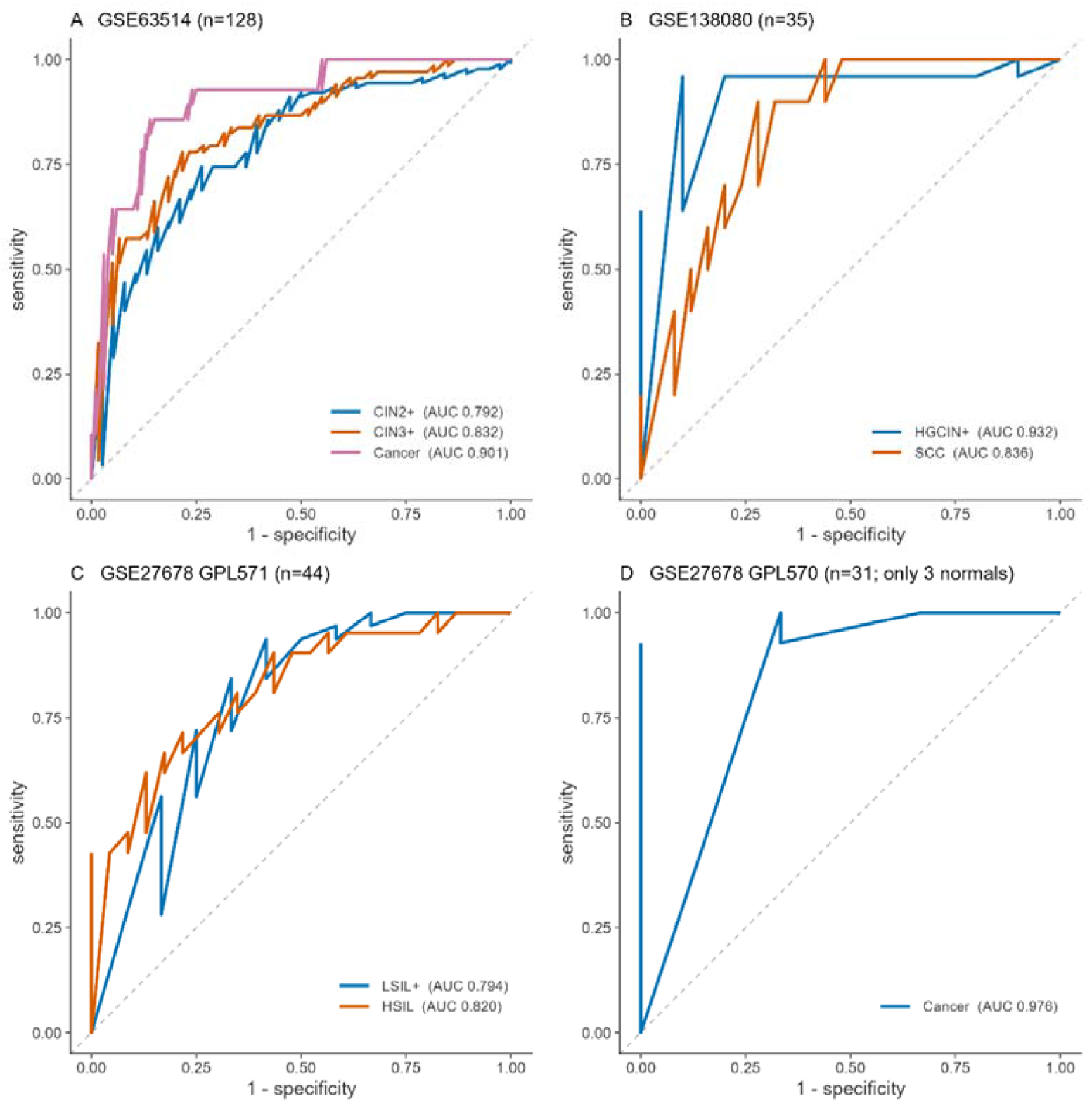
Receiver operating characteristic curves for the composite index at each pre-specified threshold, one panel per cohort, all panels on identical square axes. Areas under the curve appear in the panel legends and their confidence intervals and group sizes in Table 3. Panel D rests on three normal specimens and is reported as a binary concordance check rather than as a validation

The replication cohorts reproduce the direction but are small, and their interval estimates should be read with the group sizes attached (**Table S9b**). In GSE138080 the HGCIN or worse estimate is 0.932 (0.820 to 1.000) against 10 normal specimens, resting on 17 discordant pairs out of 250, and the SCC estimate is 0.836 (0.696 to 0.948) against 10 carcinomas. In GPL571 the LSIL or worse estimate is 0.794 (0.625 to 0.940) against 12 normal specimens and the HSIL estimate 0.820 (0.681 to 0.930). For these cohorts the trend statistic, which uses every specimen, is the more informative measure.

The GPL570 set yields 0.976 (0.905 to 1.000), but this rests on three normal specimens and two discordant pairs out of 84, so a single reclassification moves it substantially. It is reported as a binary concordance check and not as a validation. One of its three normal specimens, GSM685524, carries conflicting deposited metadata: its tissue and type fields record normal uterine cervix while its source field records squamous cell carcinoma. It was retained as normal, the conservative choice, since it generates all three discordant pairs in that cohort and its exclusion raises the estimate.

### 3.6 Pathway enrichment

Ranking the carcinoma versus normal contrast by signed significance and testing the four modules against it reproduced the module-level pattern (**Figure S3, Table S10a**). Differentiation was strongly depleted, with a normalised enrichment score of -1.889 (FDR 8.8e-05, size 7), and Synthesis was depleted at -1.638 (FDR 0.025, size 9). Neither Degradation (+0.750, FDR 0.790) nor Warburg (+0.884, FDR 0.790) reached significance.

The Synthesis result rests on two genes. PPP1R3C sits at rank 21,109 of 21,755 and GYS2 at 21,163, both above the 97th percentile of down-regulation, while the module’s remaining seven members fall between the 35th and 69th percentiles (**Table S10c**). Differentiation shows the opposite structure: all seven members occupy the top 5.1% of the ranked list, from SPRR1A at rank 20,640 to CRNN at 21,747.

The leading edge reported for each module contains only members moving in the direction of its enrichment score, so it does not list a module’s responsive genes. For Degradation, whose score is positive, the leading edge names PHKG2, AGL, PHKA1 and PHKA2 and excludes PYGB, which carries the module’s largest ranking metric at -3.062 and its 91st percentile position. Supplementary **Table S10b** therefore reports significant members by direction for each module: Synthesis 0 up and 2 down, Degradation 2 up and 1 down, Warburg 1 up and 0 down, Differentiation 0 up and 7 down. A non-significant module score for Degradation reflects absence of coordination, not absence of activity.

### 3.7 Robustness and module contribution

No exclusion set changed any conclusion (**Table S9f**). Against full-cohort estimates of 0.792, 0.832 and 0.901, removing the four specimens flagged by the pre-specified criteria gave 0.782, 0.834 and 0.907; removing the five specimens named in the earlier version of this work gave 0.800, 0.827 and 0.898; and removing GSM1551331, a post hoc exclusion reported separately, gave 0.813, 0.846 and 0.910. Every shift is smaller than the corresponding bootstrap confidence half-width of 0.086, 0.069 and 0.063. The four exclusion sets were defined by different procedures and are not combined.

The index does not depend on any single gene (**Figure S4B, Table S6)**. Removing each of the 33 genes in turn and recomputing gave a mean CIN3 or worse estimate of 0.831 with a standard deviation of 0.0045 and a range of 0.820 to 0.839, against a baseline of 0.832. The largest decrements were ESR1 at -0.0120, PHKA1 at -0.0108, CRNN at -0.0091, PPP1R3B at -0.0076 and SLC2A1 at -0.0064. The three genes undetected in normal epithelium contribute nothing: removal of PPP1R3A and of PYGM each changed the estimate by -0.0002, and removal of PHKG1 by +0.0022. Removal of PKM improved the estimate by +0.0066.

Testing each module alone as a classifier localises the signal (**Figure 2B, Table S9c**). Differentiation alone gives 0.832 (0.759 to 0.898) and Synthesis alone 0.745 (0.664 to 0.829), while Degradation gives 0.504 (0.402 to 0.604) and Warburg 0.569 (0.463 to 0.665), both intervals containing 0.5. The composite gives 0.832 (0.761 to 0.899), which is 0.0005 above Differentiation alone.

Reduced forms of the index confirm this (**Table S9d, S9e**). Across the three discovery thresholds, Differentiation alone returns 0.791, 0.832 and 0.905 against the composite’s 0.792, 0.832 and 0.901. At the highest grade of each cohort, Differentiation alone equals or exceeds the composite in three of four cohorts, the exception being the GPL570 set with three normal specimens. Combining the four modules therefore confers no discriminatory advantage over the Differentiation module alone, and no such advantage is claimed. The full panel is retained because the negative results for Degradation and Warburg are themselves findings, and those modules must be measured to be reported as unchanged.

## Discussion

Glycogen disappears from cervical epithelium as disease grade rises because the cell stops making glycogen and stops building a mature squamous epithelium, not because it starts breaking glycogen down. Across four cohorts and three specimen preparations, the Synthesis and Differentiation modules fall with grade while the Degradation and Warburg modules remain at their normal-tissue baseline. Three independent analyses agree on the negative. The Warburg module does not separate from normal epithelium at any disease grade, with p values between 0.45 and 0.62; alone it classifies CIN3 or worse at an area under the curve of 0.569 with a confidence interval spanning chance; and it shows no enrichment in the carcinoma contrast (normalised enrichment score +0.884, FDR 0.790). Degradation behaves the same way, and where its members do move they move in opposite directions, two up and one down. This is the first study to measure all four arms of cervical glycogen biology in parallel across a multi-cohort design, and the decomposition settles a question histochemistry has been unable to answer since iodine staining entered practice. Glycogen depletion is anabolic failure, not catabolic consumption. The arm the cancer-metabolism literature would have predicted is excluded, and the arm that fails is named.

The selectivity within the Synthesis module sharpens this further. Only two of its nine members change in carcinoma: GYS2, the hepatic-type glycogen synthase, and PPP1R3C, the PTG glycogen-targeting subunit of protein phosphatase 1. The ubiquitous synthase GYS1 does not move, nor do GBE1, UGP2, PGM1 or the other regulatory subunits. Glycogen storage in squamous cervical epithelium therefore appears to depend on a tissue-restricted synthetic axis rather than on the housekeeping pathway, and it is that axis which fails. The module-level enrichment result rests on these two genes and should be read as such rather than as a coordinated collapse of glycogen synthesis. The Differentiation module shows the opposite structure, with all seven members significantly reduced and all seven in the top five per cent of the ranked transcriptome, which is the signature of a programme being switched off rather than individual genes drifting.

That glycogen is lost from dysplastic cervical epithelium has been known since Schiller introduced iodine staining, and the observation remains in routine use wherever Lugol’s iodine supports visual inspection in screening programmes [12,14]. What has not been established is which part of glycogen metabolism accounts for the loss. The default expectation from the cancer metabolism literature is catabolic: a transformed cell is held to mobilise stored carbohydrate and shift toward aerobic glycolysis, so glycogen should fall because it is being consumed [23,29,45]. Our data do not support that explanation in cervical carcinogenesis. Transcript-level evidence for glycolytic activation is confined to a single transporter, SLC2A3, while GLUT1, hexokinase 2, PKM, LDHA, enolase 1, PFKL and G6PD are unchanged. Phosphorylase does not rise; PYGB falls. Where catabolic genes do move, they are phosphorylase kinase subunits and the debranching enzyme, moving modestly and inconsistently in direction.

The alternative explanation fits the biology of the tissue and of the virus. Glycogen accumulation in cervical epithelium is a property of maturation, confined to the intermediate and superficial layers of a stratified epithelium that has completed its differentiation programme[46,47]. A lesion that fails to differentiate has no compartment in which to store glycogen, so the store is never laid down rather than being drawn down. High-risk HPV provides a mechanism for exactly that failure, since the viral oncoproteins uncouple cell-cycle exit from differentiation and retain cells in a proliferative state within the suprabasal compartment [48,49]. On this account glycogen depletion is a downstream consequence of an arrested differentiation programme, and the GYS2 and PPP1R3C axis fails as part of that programme rather than as an independent metabolic event. The temporal ordering we observe is consistent with, though it does not establish, such a sequence: the Synthesis module separates from normal at CIN2 while Differentiation does not separate until CIN3.

Two implications follow. First, the continued use of iodine-based visual inspection in low-resource screening has a transcriptional rationale that is more specific than has been appreciated: the test reports a failure of squamous maturation, read through its glycogen consequence, rather than a metabolic property of malignancy. The Synthesis and Differentiation modules are the molecular counterpart of that signal, and an index built from the two performs equivalently to every other form tested, 0.825 against 0.832 for the full index with overlapping intervals, which is what a mechanistic readout requires and all it requires. This matters where cytology and HPV testing are not universally available and visual methods carry a substantial share of the screening burden [50– 52]. Second, this work does not produce a clinical test and should not be read as proposing one. ssGSEA scores are normalised within a sample set, so no cut-point derived here transfers to another cohort, let alone to a clinical specimen. The composite index is an instrument for decomposing a mechanism rather than a classifier. Every form containing the Differentiation module falls within 0.007 of every other, from 0.825 for the two-module form to 0.832 for the full index, so the discriminating information is concentrated in one module and no combination is advanced as a classifier.

The study’s strengths lie in its negative results and in how hard they were tested. The finding that catabolic and glycolytic programmes do not move is supported by three methodologically distinct analyses that could each have contradicted it. The result survives removal of any single gene, with a standard deviation across all 33 leave-one-out estimates of 0.0045, and it survives every outlier handling we tried, with each shift smaller than its own bootstrap confidence half-width. Specimens flagged by the pre-specified criteria, specimens flagged in an earlier version of this work, and two post hoc exclusions were analysed as four separate sets and never pooled. All analytical decisions, including the four taken after the data were first examined, are recorded with their rationale and deposited with the code. Reporting a signature that fails to outperform one of its own components is not an attractive result to publish, but concealing it would have left the central mechanistic claim resting on a classifier that does not work as described.

Four limitations constrain what can be concluded. No protein or metabolite data are presented, so the inference from depleted synthetic machinery to depleted glycogen remains an inference. The cell-composition confound is not resolved by this design: a specimen that has lost its superficial layer contains fewer of the cells that express differentiation genes, and convergence across three preparations is a weaker defence than it first appears, because microdissected epithelium, bulk tissue and exfoliated cytology all lose the same cell population in the same direction. Only a spatially resolved method can separate these explanations. Sample sizes outside the discovery cohort are small: ten and twelve normal specimens in the two replication cohorts, and three in the GPL570 concordance check. All four cohorts are archival series of unstated geographical origin, so these findings are not demonstrated in an African population. Three panel genes, PYGM, PHKG1 and PPP1R3A, are expressed at background in the discovery cohort, and five are absent from the GPL4133 array. One finding is sensitive to an analytical choice: median index values increase monotonically across all five discovery grades under the probe-collapse rule adopted here but reverse at the normal to CIN1 step under the alternative; that step is not significant under either, and no claim of monotonicity is made.

Glycogen depletion in cervical carcinogenesis is a failure of synthesis within a collapsing squamous differentiation programme, not catabolic activation. The loss is traceable to a tissue-restricted axis, GYS2 and PPP1R3C, while the housekeeping synthase GYS1 is retained, and the catabolic and glycolytic arms that the cancer-metabolism literature would have predicted do not move in any cohort. Three steps follow. Protein-level work must establish that GYS2 and PPP1R3C are reduced where glycogen is reduced, which requires spatially resolved methods on intact tissue and is the subject of a separate study. The temporal ordering needs a design that can resolve the two events in time rather than inferring their order from grouped cross-sectional specimens. And the signature needs testing in prospectively collected African specimens, where cervical cancer incidence and HPV genotype distribution both differ. None of this alters what the transcriptome already shows: the tissue change that visual screening detects is a differentiation failure, and the molecular target that follows from it is the differentiation programme rather than glycolysis.

## Acknowledgements

The authors thank the investigators who deposited the GSE63514, GSE138080 and GSE27678 series in the Gene Expression Omnibus. This analysis depends entirely on their data sharing.

## Ethics approval

This study analysed publicly available, de-identified gene expression data. No ethical approval or consent was required. Ethical approval and participant consent for the original specimens are reported in the primary publications of each series.

## Data availability

All data analysed are publicly available from the NCBI Gene Expression Omnibus: GSE63514 (GPL570), GSE138080 (GPL4133) and GSE27678 (GPL570 and GPL571). No new data were generated. Scripts retrieve each series directly from GEO, so no redistributed copy is required.

## Code availability

The complete analysis pipeline is available at https://github.com/oyedele-ajayi/cervical-glycogen-anabolic-failure and archived at Zenodo (DOI: 10.5281/zenodo.20289069), which resolves to the current version. The repository contains every script in execution order, a documented run order, a decision record of all analytical choices, and a verification script that checks each value reported here against the analysis output.

## Supplementary information

The supplementary workbook contains the locked 33-gene panel and its per-cohort coverage, full differential expression results, module and index scores for every specimen, trend and discrimination statistics, the robustness analyses, and the decision record. Supplementary Figures S1 to S4 are provided separately.

## Declaration of generative AI and AI-assisted technologies in the writing process

During the preparation of this work the authors used Claude (Anthropic) to assist with drafting, structuring and language editing of the manuscript, and with developing and documenting the analysis code. The authors reviewed and edited all content, verified all reported values against the deposited analysis output, and takes full responsibility for the content of the publication.

## Author contributions

**Oyedele O. Ajayi:** Conceptualization, Methodology, Software, Formal analysis, Project administration, Writing – original draft, Writing – review and editing.

**Efosa B. Odigie:** Supervision, Data curation, Writing – review and editing. Identified, screened and retrieved the transcriptomic datasets from public repositories

**Samson O. Adesite:** Investigation, Formal analysis, Writing – review and editing. Curated the glycogen metabolism gene set, performed functional enrichment analysis and contributed biological interpretation.

**Theophilus O. Erameh:** Validation, Formal analysis, Writing – review and editing. Conducted statistical validation, sensitivity analyses and reproducibility assessment.

**Anietie M. Archibong:** Investigation, Writing – review and editing. Contributed cervical pathology expertise and interpretation of the findings in relation to HPV-associated metabolic change and visual inspection screening.

**Uchechuckwu B. Eziagu:** Validation, Visualization, Writing – original draft, Writing – review and editing. Performed external dataset validation and prepared the figures.

