## Supplementary Figures for "ANABOLIC FAILURE, NOT CATABOLIC ACTIVATION, UNDERLIES GLYCOGEN DEPLETION IN CERVICAL CARCINOGENESIS: A MULTI-COHORT TRANSCRIPTOMIC ANALYSIS"

**Figure S1a: GSE63514, discovery cohort, n = 128**

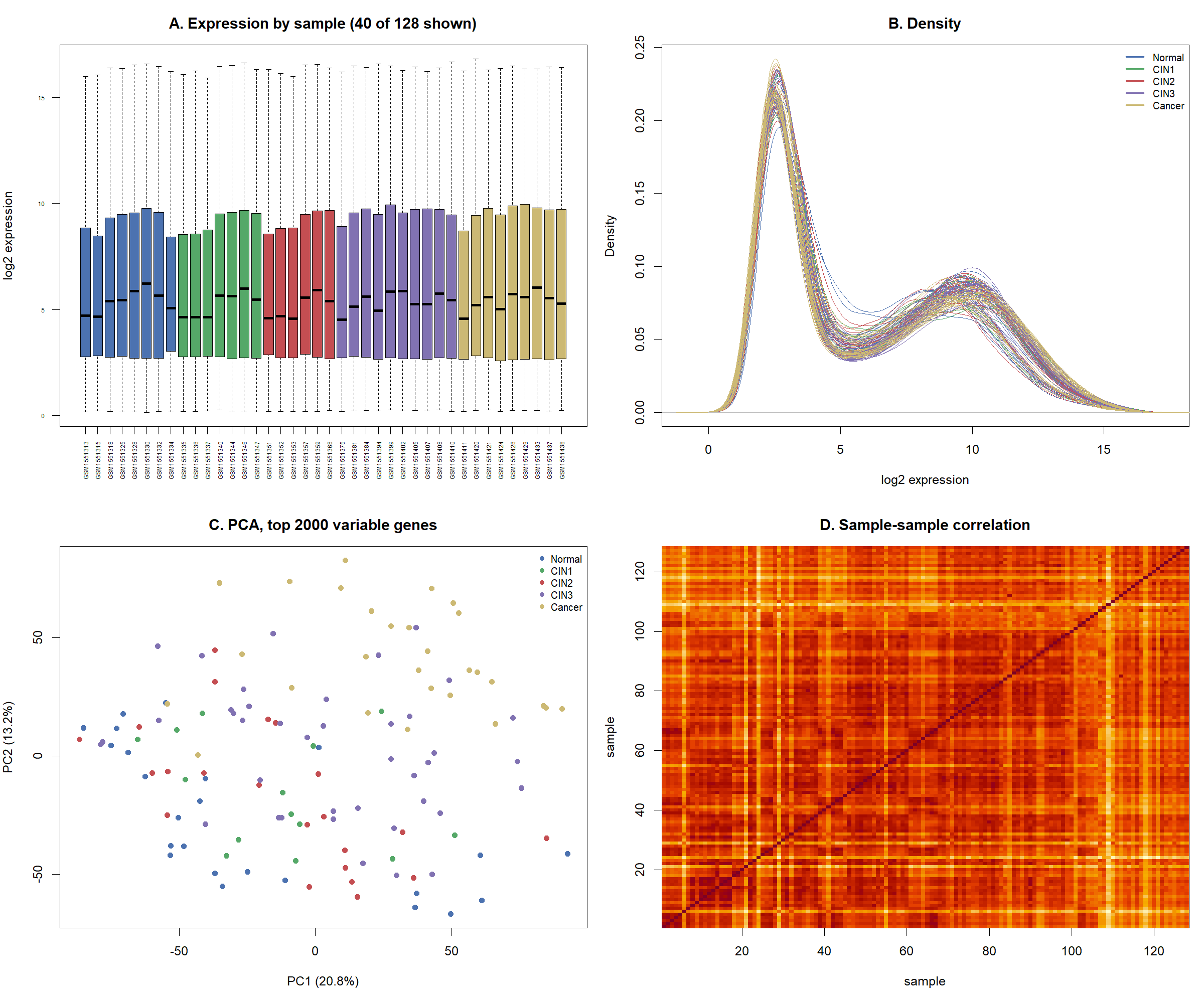

**Figure S1b: GSE138080 n = 35**

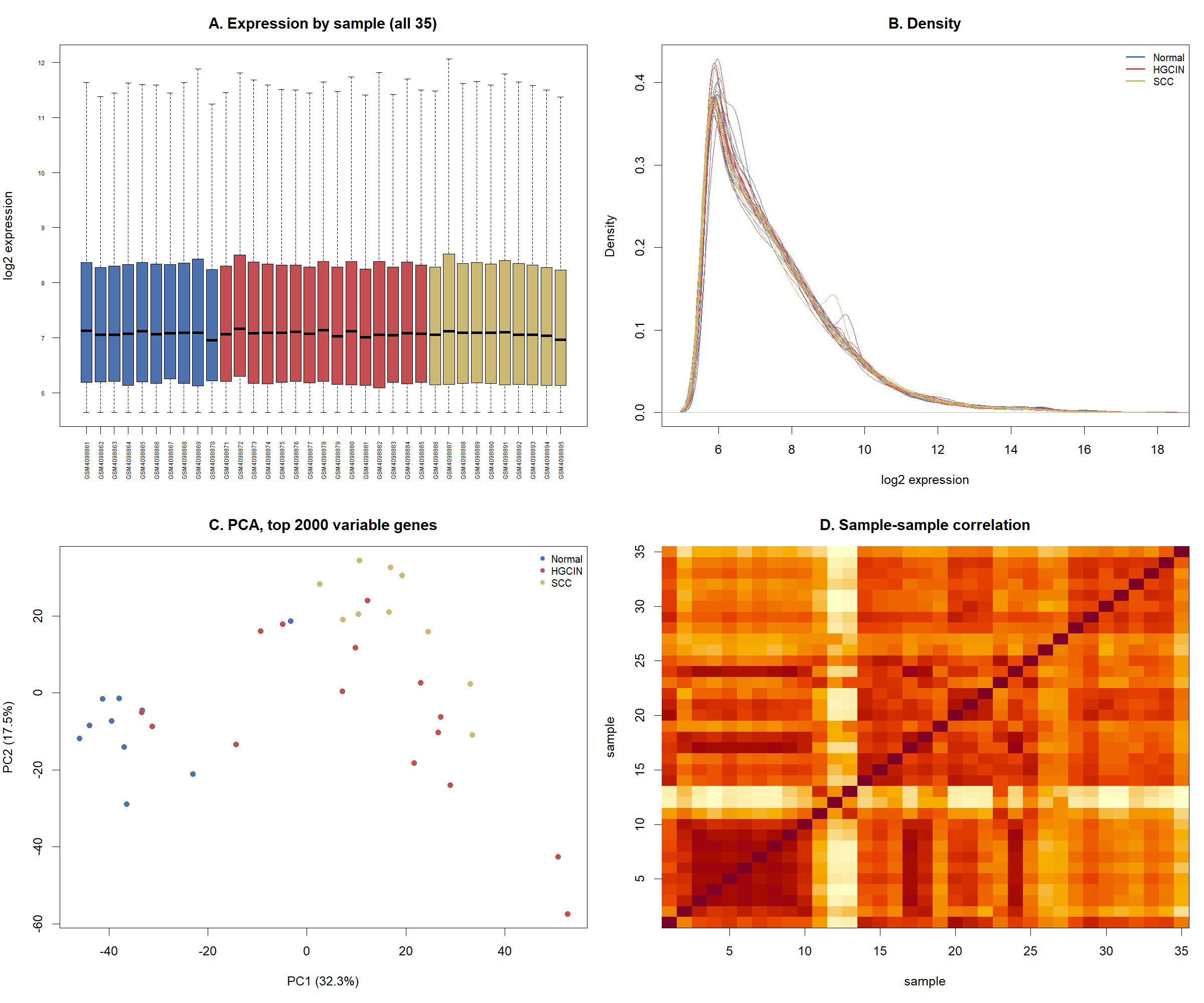

**Figure S1c: GSE27678 GPL571 n = 44**

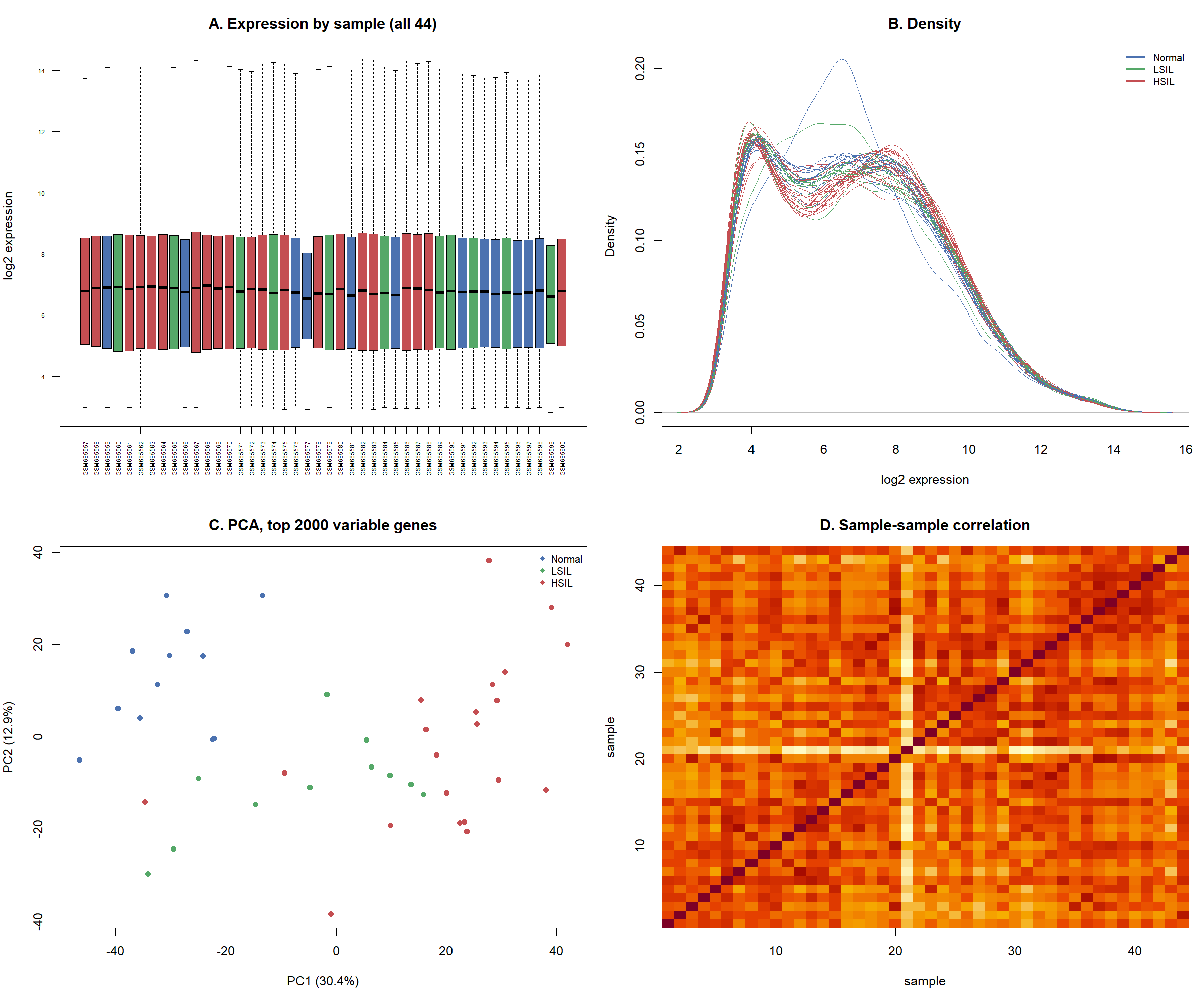

**Figure S1d: GSE27678 GPL570 n = 31**

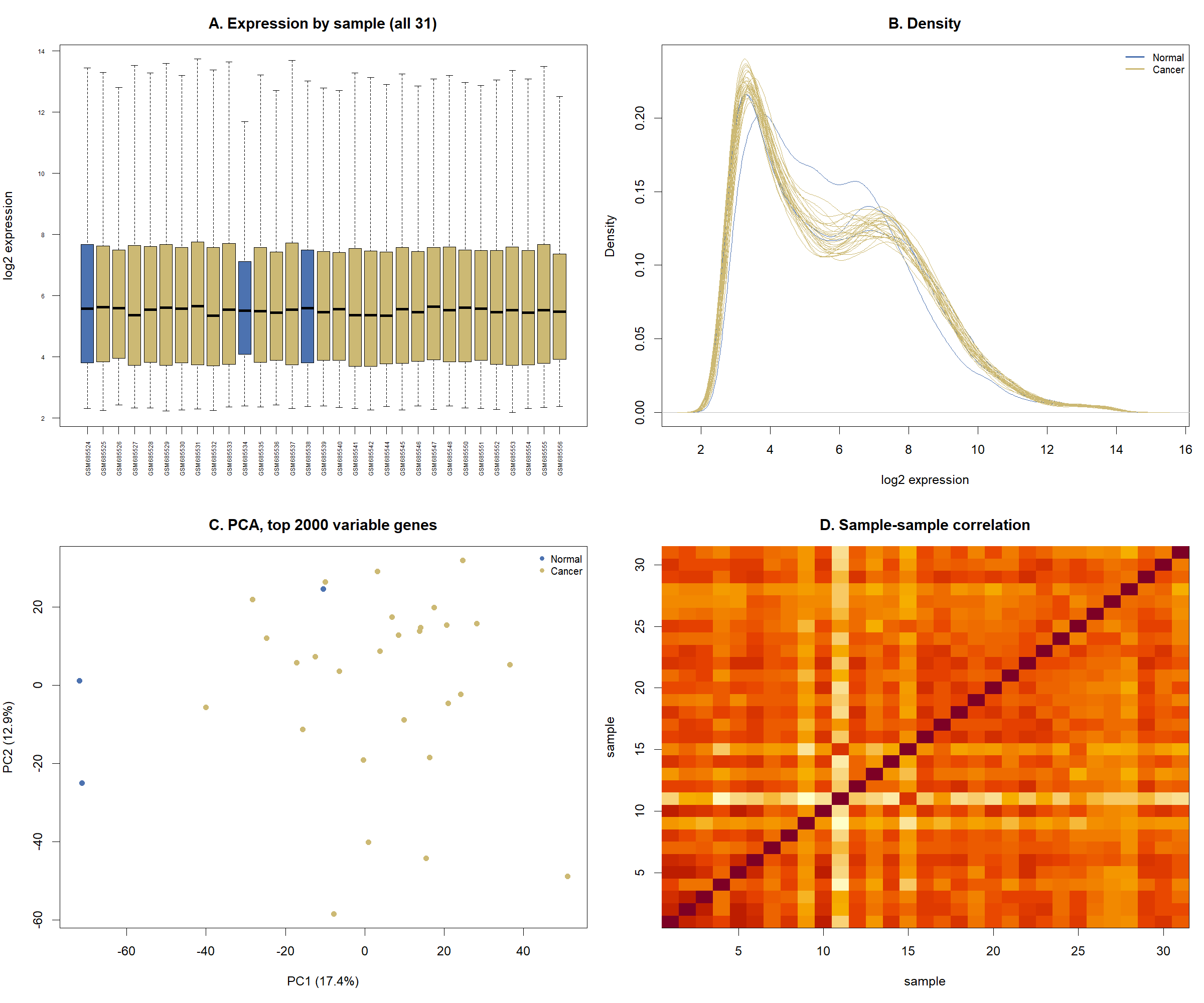

Figure S1. Quality control of the four expression cohorts*.*

*Each page shows one cohort: S1a, GSE63514 discovery cohort, laser-capture-microdissected epithelium, n = 128; S1b, GSE138080, bulk tissue, n = 35; S1c, GSE27678 GPL571, exfoliated cytology, n = 44; S1d, GSE27678 GPL570, exfoliated cytology, n = 31. Panels are identical across cohorts. (A) Distribution of log2 expression per sample. For the discovery cohort, 40 of the 128 samples are shown, drawn at random with a fixed seed, because 128 boxes are illegible at page width; all other cohorts show every sample. (B) Density of log2 expression, one curve per sample. (C) Principal component analysis of the 2000 most variable genes, showing PC1 against PC2 with the percentage of variance each explains. (D) Pearson correlation between every pair of samples, ordered as in the expression matrix, with darker colour indicating higher correlation. Points and curves are coloured by disease grade throughout: Normal, CIN1, CIN2, CIN3 and Cancer in the discovery cohort; Normal, HGCIN and SCC in GSE138080; Normal, LSIL and HSIL in GPL571; Normal and Cancer in GPL570. CIN, cervical intraepithelial neoplasia; HGCIN, high-grade cervical intraepithelial neoplasia; LSIL and HSIL, low-grade and high-grade squamous intraepithelial lesion; SCC, squamous cell carcinoma.*

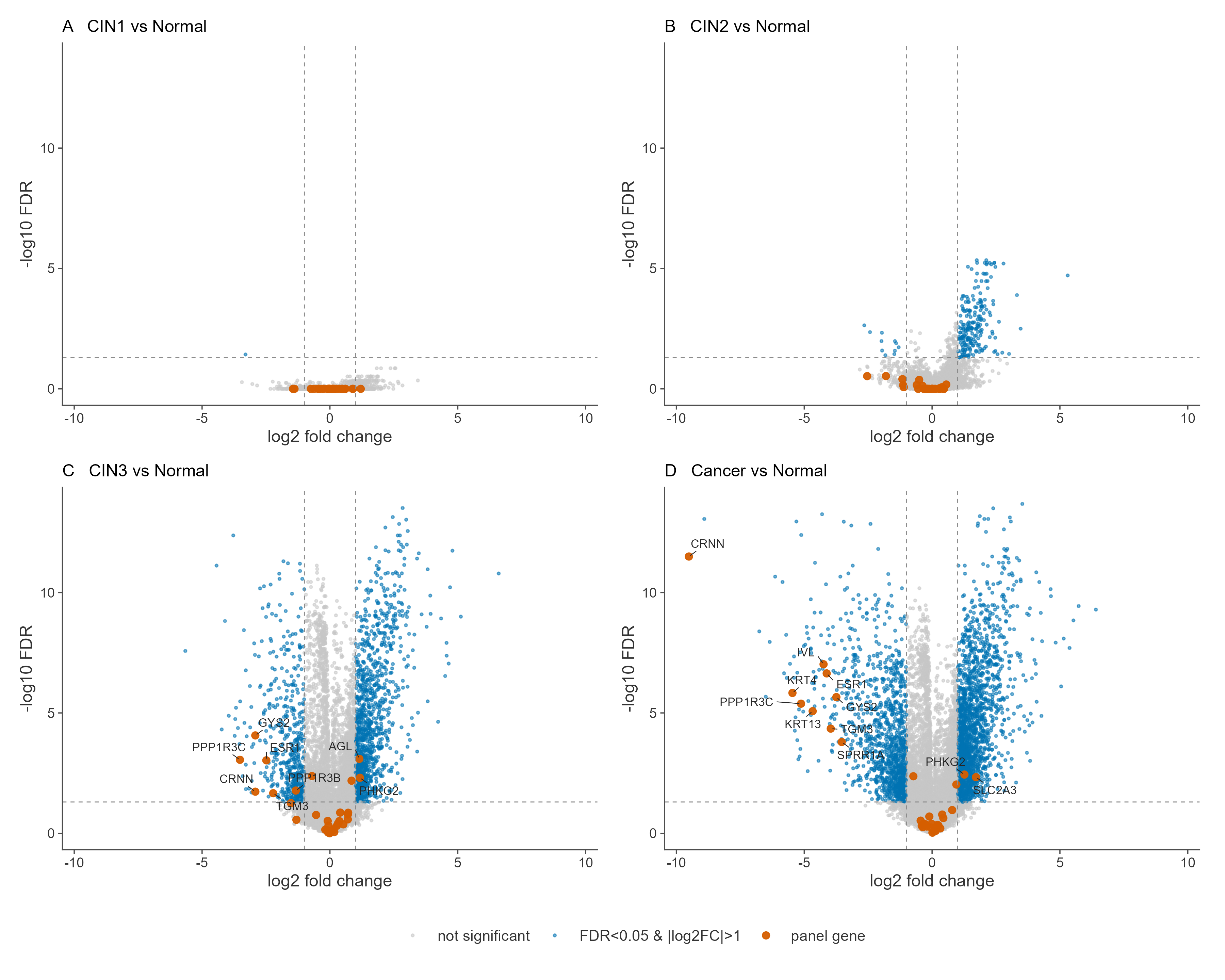

**Figure S2. Transcriptome-wide differential expression in the discovery cohort.**

*Volcano plots for four contrasts against histologically normal epithelium in GSE63514 (n = 128): (A) CIN1, (B) CIN2, (C) CIN3, (D) invasive cancer. Each point is one gene, plotted as log2 fold change against the negative log10 of the Benjamini-Hochberg adjusted p value. Grey points are not significant. Coloured points meet both FDR below 0.05 and absolute log2 fold change above 1. The 33 panel genes are overplotted in a contrasting colour at larger size and labelled where they meet both thresholds. Dashed lines mark FDR 0.05 and log2 fold change of plus and minus 1. All four panels share identical axes so magnitudes are directly comparable. FDR, false discovery rate; CIN, cervical intraepithelial neoplasia*

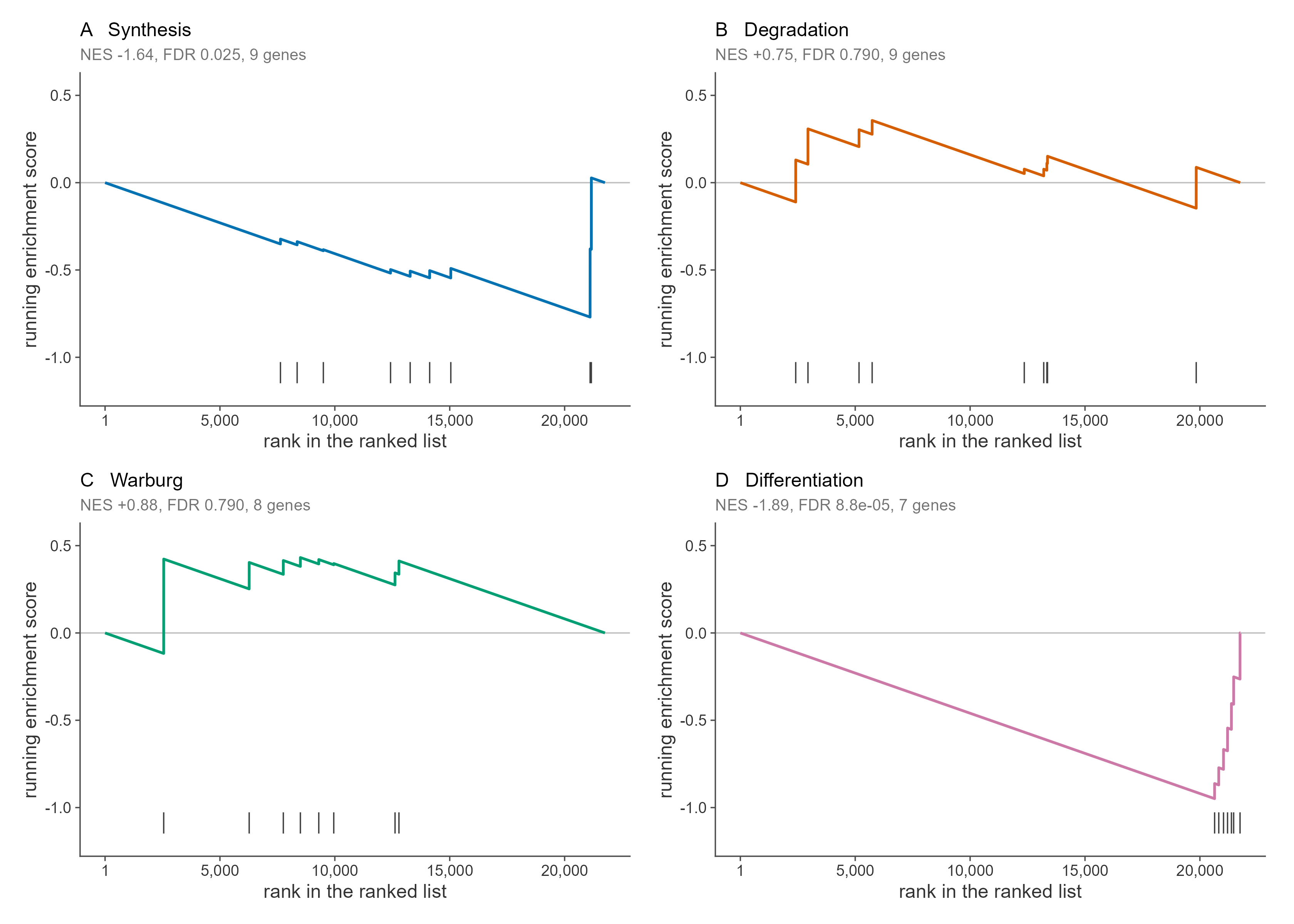

**Figure S3. Running enrichment of the four metabolic modules.**

*Gene set enrichment analysis of each module against the transcriptome-wide ranked list for invasive cancer versus normal epithelium in GSE63514 (n = 128), ranked by the moderated t statistic. (A) Synthesis, (B) Degradation, (C) Warburg, (D) Differentiation. The curve is the weighted running enrichment score as each gene is encountered in rank order. Tick marks below each curve mark the positions of that module’s member genes in the ranked list. Normalised enrichment score, false discovery rate and module size are given in each panel subtitle. All four panels share the same vertical scale, so curve heights are comparable between modules. A curve that departs downward indicates coordinate loss of expression in cancer; one that stays near zero indicates no coordinate shift.*

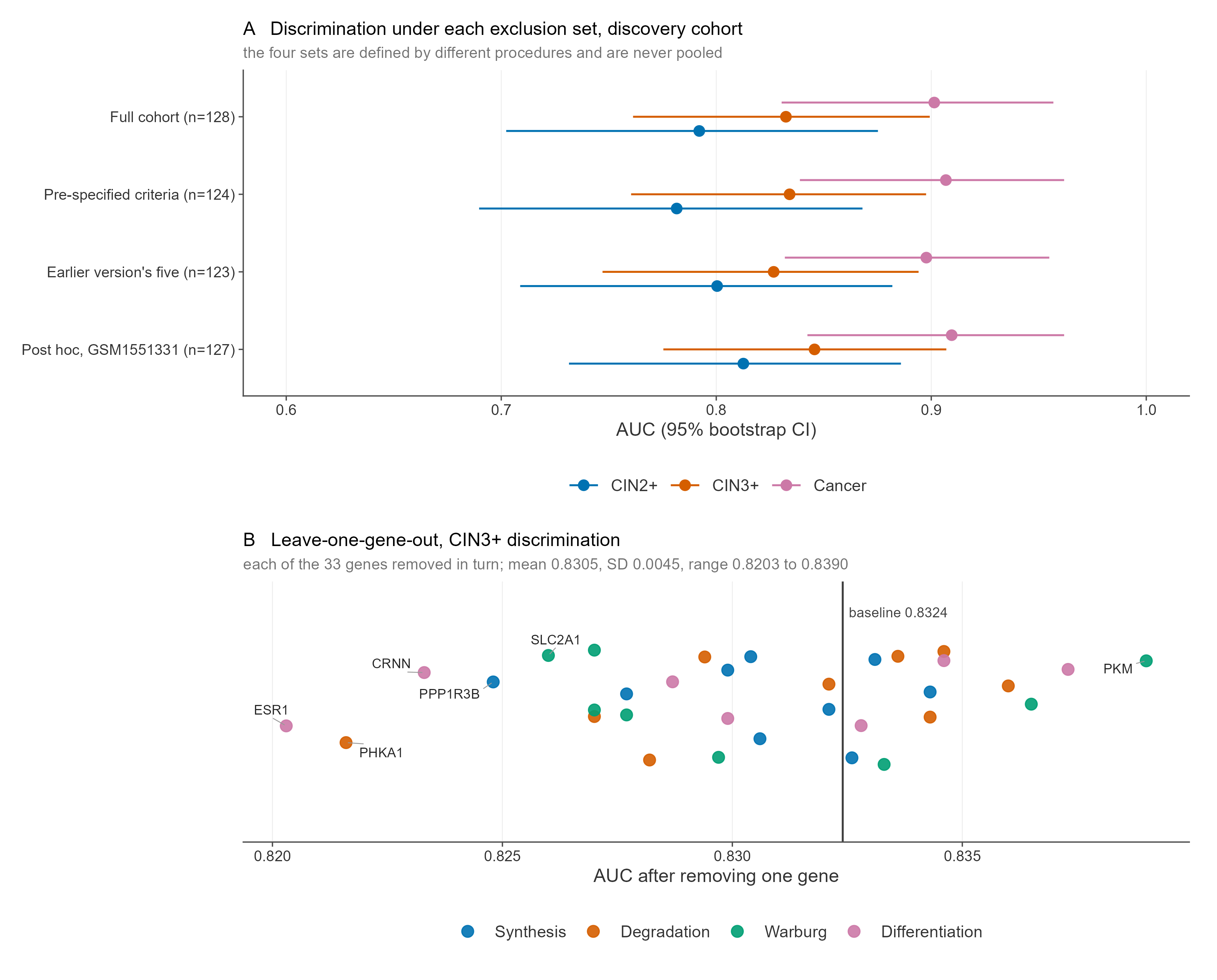

**Figure S4. Robustness of the discovery estimate.**

(A) *Discrimination of the glycogen depletion index under four sample exclusion sets, each defined by a different procedure and never pooled: the full cohort (n = 128), the pre-specified outlier criteria (n = 124), the five samples named in an earlier version of this work which meet neither stated criterion (n = 123), and a post hoc exclusion of GSM1551331 (n = 127). Each point is the area under the receiver operating characteristic curve with its 95% bootstrap confidence interval from 2000 stratified resamples, shown at three thresholds: CIN2 or worse, CIN3 or worse, and invasive cancer****.*** *(B) Leave-one-gene-out analysis of CIN3 or worse discrimination. Each point is the area under the curve recomputed with one of the 33 panel genes removed, coloured by module. The vertical line is the baseline estimate using all 33 genes. The five genes whose removal lowers the estimate most, and the one whose removal raises it most, are labelled. Mean, standard deviation and range across all 33 removals are given in the panel subtitle. AUC, area under the receiver operating characteristic curve; CIN, cervical intraepithelial neoplasia*
